# A low-cost and versatile paramagnetic bead DNA extraction method for *Mycobacterium ulcerans* environmental surveillance

**DOI:** 10.1101/2024.05.31.596290

**Authors:** Jean Y. H. Lee, Jessica L. Porter, Emma C. Hobbs, Pam Whiteley, Andrew H. Buultjens, Timothy P. Stinear

**Author notes:** Address correspondence to Timothy P. Stinear.

## Abstract

In Australia, native possums are an animal reservoir for *Mycobacterium ulcerans*, the causative agent of Buruli ulcer, a neglected tropical skin disease that can progress to extensive ulceration with deformity and disability. Surveillance of possum excreta for shedding of *M. ulcerans* can be used to inform geospatial modelling to predict locations at which humans are at increased risk of disease acquisition. This discovery provides opportunities to disrupt transmission pathways. However, the significant expense of commercial kits used for DNA extraction from environmental samples in large scale surveillance studies can hinder implementation of public health measures. To address this barrier, we developed a low-cost method for extraction of nucleic acids from possum excreta, possum tissue swabs, and mycobacterial cultures, using a guanidinium thiocyanate lysis solution and paramagnetic beads for DNA clean-up. In 96-well plate format for high-throughput processing, the SPRI-bead DNA extraction method for possum excreta was 3-fold less sensitive but only 1/6 the cost of a widely used commercial kit. While a SPRI-bead microtube-based extraction method for tissue swabs sampled from possums was 4-fold more sensitive and 1/5 the cost of the corresponding commercial kit. Furthermore, when used for preparing DNA from pure mycobacterial cultures, the SPRI-bead method produced genomic DNA with quality metrics comparable to more laborious techniques. The methods described here provide an economical means to continue large-scale *M. ulcerans* environmental surveillance that should facilitate efforts to halt the spread of Buruli ulcer in Victoria, Australia, with potential for applicability in other endemic countries.

**IMPORTANCE:** Buruli ulcer is a neglected tropical skin disease, with an incidence that has dramatically increased in temperate southeastern Australia over the last decade. In this region of the world Buruli ulcer is a zoonosis, Australian native possums are a major wildlife reservoir of the causative agent, *Mycobacterium ulcerans*, and mosquitoes the vector to humans. Infected possums shed *M. ulcerans* in their excreta, and excreta surveys using PCR to screen for the presence of pathogen DNA are a powerful means to predict future areas of Buruli ulcer risk for humans. However, excreta surveys across large geographic areas require testing of many thousands of samples. The cost of commercial DNA extraction reagents used for preparing samples for PCR testing can become prohibitive to effective surveillance. Here, we describe a simple, low-cost method for extracting DNA from possum excreta using paramagnetic beads. The method is versatile and adaptable to a variety of other sample types including swabs collected from possum tissues and pure cultures of mycobacteria.

## INTRODUCTION

Buruli ulcer (BU) is a chronic bacterial infection affecting subcutaneous tissues, caused by *Mycobacterium ulcerans*. Reported in 33 countries across Africa, the Americas, Asia and the Western Pacific, BU is classified by the World Health Organization as a neglected tropical disease [1, 2, 3]. Although Sub-Saharan Africa has the highest global burden of disease, of the countries that report annual suspected case numbers, Australia was second only to the Democratic Republic of the Congo in 2022, with 343 and 558 new reported cases, respectively [4, 5]. In temperate south-eastern Australia, over the last 26 years the endemic region for BU has expanded from selected coastal regions in the state of Victoria, to the major metropolitan centres of Geelong then Melbourne, with state-wide case numbers escalating to 362 notifications in 2023, the highest since the disease became notifiable to the Victorian Department of Health in 2004 [6, 7]. Extensive surveillance studies in this region have led to the discovery that BU is a zoonotic infection, with Australian native possums a wildlife reservoir of *M. ulcerans* and mosquitoes a primary vector for transmission to humans [8, 9, 10, 11, 12, 13]. In Africa, a *M. ulcerans* wildlife reservoir analogous to possums is yet to be discovered, and while there is a lack of evidence implicating mosquitos as a vector, local studies suggest a role for aquatic insects spreading the bacterium [14, 15, 16, 17].

While *M. ulcerans* infection has been reported in both native Australian [11, 13, 18, 19, 20], and domesticated animal species both in Australia and overseas [1, 19, 21, 22, 23], two possum species, the common ringtail (*Pseudocheirus peregrinus*) and common brushtail (*Trichosurus vulpecula*) possums have been identified as major reservoirs for *M. ulcerans* infection in Victoria, Australia [9, 10, 11, 12, 13]. The percentage of affected possums varies by region. In the endemic area of Point Lonsdale, a 2010 study reported 38% (N = 16/42) of ringtail possums and 24% (N = 5/21) of brushtail possums to have either laboratory-confirmed *M. ulcerans* lesions or *M. ulcerans* PCR-positive excreta [11]. This study observed that *M. ulcerans* DNA was detected in 41% of possum faecal samples collected in a Buruli endemic area compared to less than 1% of faecal samples collected from non-endemic areas (p<0.0001) [11]. A 2014 study performed in the Mornington Peninsula, detected *M. ulcerans* in 9.3% (N = 20/216) of ringtail and 66.6% (N = 4/6) of brushtail possum excreta sampled [9]. This study also demonstrated a statistically significant (p<0.0001), non-random clustering of *M. ulcerans* positive possum excreta within a 0.42 km radius of the addresses of confirmed human cases of BU [9]. The hypothesis that possums are a sentinel species that can be used to predict human BU cases prompted more sophisticated confirmatory studies [8, 10]. Culminating in the 2024 study by Mee et al., that employed spatial scanning statistics to show an overlap between *Aedes notoscriptus* mosquitos that tested positive for *M. ulcerans*, with *M. ulcerans* positive possum excreta, and human BU cases; paired with metabarcoding analyses that demonstrated the feeding of individual mosquitos on both possums and human; and targeted enrichment genome capture to link the transmission chain between mosquitos, native possums and humans [10].

Despite recent advances in understanding the transmission of *M. ulcerans*, studies of the environmental ecology of this pathogen remain challenging, primarily due to its slow growth and concomitant difficulties isolating this organism in pure culture from microbially complex environmental samples [11, 24]. A TaqMan quantitative polymerase chain reaction (qPCR) targeting IS*2404* has been the main methodology employed to infer the presence of *M. ulcerans* in the environment [8, 10, 11, 12, 13]. The IS*2404* insertion sequence encodes a 327 amino acid transposase [25] that recurs approximately 200 times in the *M. ulcerans* genome [25, 26, 27], making it an ideal target for PCR amplification. Prior to testing, possum excreta samples require DNA extraction, the cost of commercial DNA extraction kits can become prohibitively expensive in large scale surveillance studies involving thousands of samples. To overcome this barrier, we sought to develop a significantly lower-cost DNA extraction method.

Australian brushtail and ringtail possums are predominantly folivores, with leaves of *Eucalyptus* species being their primary food source, supplemented by shrub foliage, flowers and fruit; and in the case of brushtail possums, small quantities of animal material [28, 29]. Subsequently, possum excreta are rich in plant matter [29], including humic substances, which are compounds resulting from the decomposition of organic matter [30]. Due to similar chemical properties to nucleic acids, humic compounds are typically co-purified during standard nucleic acid extraction methods, these compounds are established inhibitors of PCR reactions [31] and most other subsequent analyses performed on DNA [32, 33]. Therefore, additional treatment is required to remove humic substances when performing DNA extraction of material that is known to be rich in organic matter. Chemical flocculation, which employs multivalent cations (such as Al3+) to bind and remove organic inhibitors has been demonstrated as an effective treatment for the removal of humic compounds when extracting DNA from environmental samples such as soil [32, 34, 35].

We have previously demonstrated the utility of solid-phase reversible immobilisation (SPRI) paramagnetic beads as a low-cost method for the purification of nucleic acids [36]. Here we show that using a pre-treatment with aluminium ammonium sulphate (AlNH_4_(SO4)_2_, also known as alum), combined with mechanical and chemical lysis with bead beating and a guanidinium thiocyanate (GITC) based lysis solution, followed by nucleic acid extraction with SPRI beads, we can obtain *M. ulcerans* DNA from possum excreta without comprising diagnostic yield, at 1/6 the cost of commercial reagents. Furthermore, the method was demonstrated to be adaptable for DNA extraction from biological swabs collected from possums with suspected *M. ulcerans* infection and for the extraction of mycobacterial genomic DNA.

## MATERIALS AND METHODS

### Biological specimen collection

Voided possum excreta included samples collected as part of a structured 12 month possum excreta surveillance program spanning across 350 km2 in the Mornington Peninsula, 70 km south of Melbourne, conducted during two seasonal periods: a ‘summer survey’ from December 19, 2018 to March 14, 2019 and ‘winter survey’ from May 28 to September 19, 2019 [8]. Additional samples were collected as part of subsequent surveillance conducted using the same standardised manner described in Vandelanoote et al., [8] extending to inner metropolitan Melbourne, collected over 2021 to 2023. Excreta were stored at -20oC until time of processing. Dry, flocked swabs (Copan 8155C1S) were used to sample cutaneous lesions, and in the case of necropsied possums, organs and/or body cavities (oral, cloaca, pouch) from wild, native possums during active trap-and-release and passive necropsy-based surveillance studies conducted around Melbourne and Geelong, Victoria, between November 2021 and December 2022 (Hobbs et al., 2024, manuscript under review). Collected swabs were stored in individual tubes at -20oC until time of processing. *M. ulcerans* clinical isolate JKD8049 (or DNA from this isolate) was used as positive control material for all experiments [26].

### Preparation of contrived M. ulcerans-positive possum excreta samples

Possum excreta spiked with a standardised inoculum of *M. ulcerans* was made as follows: 20 mg of *M. ulcerans* JKD8049 harvested from a Brown and Buckle slope was added to 5 ml of nuclease-free water in a 50 ml centrifuge tube. To create a homogenous suspension, a sterile 1 ml pipette tip was added and the tube vortexed. Five grams of possum excreta known to be *M. ulcerans* IS*2404* qPCR negative, crushed to the consistency of filter coffee grounds, was added to the culture and pipette tip, then vortexed to mix. The sample was stored at 4oC and allowed to soak overnight before testing.

### DNA extraction buffers, solutions, and paramagnetic beads

The 15 mM AlNH_4_(SO4)_2_ wash buffer contained 15 mM AlNH_4_(SO4)_2_.12H_2_O and 100 mM Tris HCl (pH 6.0). The 6M GITC lysis buffer comprised of 6 M GITC, 150 mM Tris HCl (pH 8.0), 90 mM dithiothreitol, 30 mM ethylenediaminetetraacetic acid (pH 8.0), 3% Triton-X 100 (volume/volume) and was stored protected from light. Note that this solution precipitates when stored below 28oC. Solid-phase reversible immobilisation (SPRI) on carboxylated paramagnetic beads (SpeedBeed Magnetic Carboxylate Modified Particles 100 ml, Azide 0.05% 65152105050350, GE Healthcare) were functionalised for DNA-binding as described by [37] and stored at 4oC, protected from light. Unless otherwise specified, chemicals were purchased from Sigma Aldrich.

### Possum excreta DNA extraction using SPRI beads

Up to 100 mg of possum excreta crushed to the consistency of filtered coffee grounds was added to a pre-autoclaved (121oC, 15 minutes) screw cap tube containing 1 g of 0.1 mm zirconium oxide beads. A 1 ml aliquot of 15 mM AlNH_4_(SO4)_2_ wash solution was added, the tube vortexed for 1 minute, then centrifuged at 17,000 x g for 1 minute, and 700 µl of the supernatant discarded. A 700 µl aliquot of 6M GITC lysis solution was added, and the tube subjected to 1 cycle of 6000 rpm for 40 seconds in a bead beater. The tube was centrifuged at 17,000 x g for 1 minute and 600 µl of supernatant was transferred to a new microcentrifuge tube. To the supernatant, 225 µl of 10 M ammonium acetate, then 75 µl of 20% (weight/volume) sodium docecyl sulfate were added, with inversion of the tube following the addition of each. The tube was centrifuged at 17,000 x g for 1 minute and 300 µl of the supernatant transferred to a new microcentrifuge tube and 200 µl of 100% ethanol added.

For each sample to be processed, an 85 µl aliquot of functionalised DNA-binding SPRI beads was added into a well of a 96-well plate, then 165 µl of the supernatant/ethanol mix was added to the same well, with gentle mixing by pipetting. After all samples and controls were aliquoted, the 96-well plate was incubated at room temperature for 5 minutes to allow SPRI-bead and DNA binding. The plate was then mounted on a magnetic rack until the supernatant was clear, with the DNA-SPRI bead complexes fixed to the bottom of the plate by the magnets. The supernatant was discarded by pipetting, with the plate still mounted on the magnetic rack. DNA-SPRI bead pellets were resuspended in 250 µl of 80% (volume/volume) ethanol with the plate off the magnetic rack, then placed back on the magnet to discard the supernatant, and the wash repeated a second time. Removal of the plate and resuspension of the DNA-SPRI bead pellets was required with each wash to remove any residual contamination. Pellets were air-dried at room temperature for 5 minutes (or until matte in appearance, avoiding a cracked pellet which indicates over drying and is associated with low DNA yield) with the plate on the magnet. The plate was then removed from the magnetic rack and DNA eluted from the SPRI beads with a 1-minute incubation in 30 µl of nuclease-free water. The plate was remounted on the magnetic rack and the eluate transferred to a clean 96-well plate then stored at -20oC if not immediately tested.

### Possum excreta DNA extraction using commercial kit

The DNeasy PowerSoil Pro Kit (Qiagen Cat 47016) was used as per manufacturer’s instructions.

### Swabs collected from possums

Swabs were inoculated into 1 ml of phosphate buffered saline (PBS) and vortexed for 1 minute, 200 µl aliquots of the sample solution were processed by either the in-house method or using the DNeasy Blood & Tissue Kit (Qiagen Cat 69504) per manufacturers’ instructions.

For the SPRI-bead method, the 200 µl sample aliquot was inoculated directly into a pre-autoclaved, screw cap tube containing 1 g of 0.1 mm zirconium oxide beads and 500 µl of 6M GITC lysis solution. The tube was subjected to 1 cycle of 6000 rpm for 40 seconds in a bead beater then processed as outlined for possum excreta extraction, up to and including the DNA precipitation step. To the 500 µl DNA/ethanol mix in a 1.5 ml microcentrifuge tube, 255 µl of functionalised DNA-binding SPRI beads was added and mixed by gentle pipetting. The tube was incubated for 5 minutes at room temperature, then placed in a 1.5 ml microcentrifuge magnetic rack until the supernatant was clear. The following steps were performed with the tube mounted in the magnetic rack without removal: the supernatant was discarded; the DNA-SPRI bead pellet was washed twice (without resuspending) with 1.5 ml of 80% (volume/volume) ethanol and the supernatant discarded after each wash; The pellet was airdried at room temperature for 5 minutes. The tube was then removed from the magnetic rack and DNA eluted from the SPRI beads with a 1-minute incubation in 100 µl of nuclease-free water. The tube was remounted in the magnetic rack and the eluate transferred to a clean 1.5 ml microcentrifuge tube then stored at -20oC if not immediately tested.

### DNA extraction from mycobacterial culture

A heaped 10 µl loop of JKD8049 *M. ulcerans* harvested from a Brown and Buckle slope was inoculated directly into a pre-autoclaved, screw cap tube containing 1 g of 0.1 mm zirconium oxide beads and 700 µl of 6M GITC lysis solution. The sample was processed from the bead beating step as outlined above, until DNA precipitation with ethanol, when 600 µl of supernatant was transferred to a new microcentrifuge tube then 400 µl of 100% ethanol was added, and the tube inverted to mix. The 1 ml of supernatant/ethanol mix was evenly divided between 2 microcentrifuge tubes and 255 µl of functionalised DNA-binding SPRI beads were added to each tube. Tubes were incubated for 5 minutes at room temperature, then placed in a 1.5 ml microcentrifuge magnetic rack until supernatants were clear. The remaining extraction process of ethanol washes and DNA elution in 1.5 ml tubes were as outlined for the possum swab extraction, except following elution in 100 µl of nuclease-free water the tubes were remounted in the magnetic rack and both eluates were pooled together in a clean microcentrifuge tube and stored at -20oC until required for testing. Genomic DNA was quantified using a fluorometer (Quibit 4 Fluorometer, Thermo Fischer), and quality was assessed with the 4200 TapeStation System (Agilent).

### IS2404 TaqMan quantitative PCR

Adapted from the method previously described, using primers targeting the IS*2404* insertion sequence in *M. ulcerans* [20], with TaqMan Exogenous Internal Positive Control Reagents - VIC Probe (Applied Biosystems Cat # 4308323) and SensiFAST probe No-ROX Kit (Bioline Cat # BIO-86005), each 20 µl reaction consisted of: 2x SensiFAST Probe No-ROX mix, 0.4 µM IS*2404* TF, 0.4 µM IS*2404* TR, 0.1 µM IS*2404* TP, 10x TaqMan Exo IPC Mix, 50x TaqMan Exo IPC DNA, 3.2 µl of nuclease free water and 2 µl of sample DNA. No-template controls, *M. ulcerans* genomic DNA positive control, and TaqMan internal positive control block (indicates if PCR inhibition has occurred) were included in every qPCR run. Amplification and detection were performed on the QuantStudio 1 (Thermo Fischer) platform using the following program: 95oC for 5 minutes, then 45 cycles of 95oC for 10 seconds and 60oC for 20 seconds. A cycle threshold (Ct) <=40 was interpreted as positive for the presence of *M. ulcerans*. This cut-off was in accordance with previous performance evaluations of the assay [8, 38]. Oligonucleotides were purchased from Integrated DNA Technologies. Run analyses were performed using Design & Analysis Software v2.5.0 (Thermo Fischer Scientific).

### Statistical analyses

Data comparisons were performed using GraphPad Prism (v 10.2.0).

## RESULTS

### Development of a SPRI-bead DNA extraction and purification method for possum excreta specimens

An overview of the extraction method developed is shown in Figure 1A. Various measures were employed to reduce the quantity of humic compounds present in the final DNA extractions, that would interfere with subsequent PCR analyses. Possum excreta were crushed to a fine ground coffee consistency to ensure older samples that were solidified from prolonged environmental exposure would adequately absorb the 15 mM AlNH_4_(SO4)_2_ (pH 6.0) wash solution. At this pH, Al3+complexes with humic substances and precipitates [35, 39]. After centrifugation, the washed pellet was incubated with the 6M GITC lysis solution (pH 8.0) and subjected to bead-beating to ensure efficient mycobacterial cell wall disruption [40]. This change in pH to 8.0, precipitates superfluous Al3+ preventing unwanted complexing of DNA [32, 35] released during bead beating. The lysate was further clarified with the addition of ammonium acetate and sodium dodecyl sulphate to precipitate proteinaceous materials. Digestion of possum excreta in the 6M GITC lysis solution for more than 45 minutes prior to protein precipitation should be avoided as it results in the release of excessive humic substances and subsequent PCR inhibition. While the method does not specify an incubation step between the addition of the 6M GITC lysis solution and protein precipitation, extended digestion may be an issue when processing many samples. After centrifugation, one third of the resulting lysate was mixed with ethanol, and approximately one third of the DNA/ethanol mix was transferred to a 96-well plate containing SPRI-beads, for a final SPRI:DNA ratio of 0.5. The DNA-SPRI bead complex was immobilised using a magnet, washed with ethanol twice to remove impurities and then the DNA eluted from the beads using a small volume (30 µl) of nuclease-free water.

**FIG 1.**
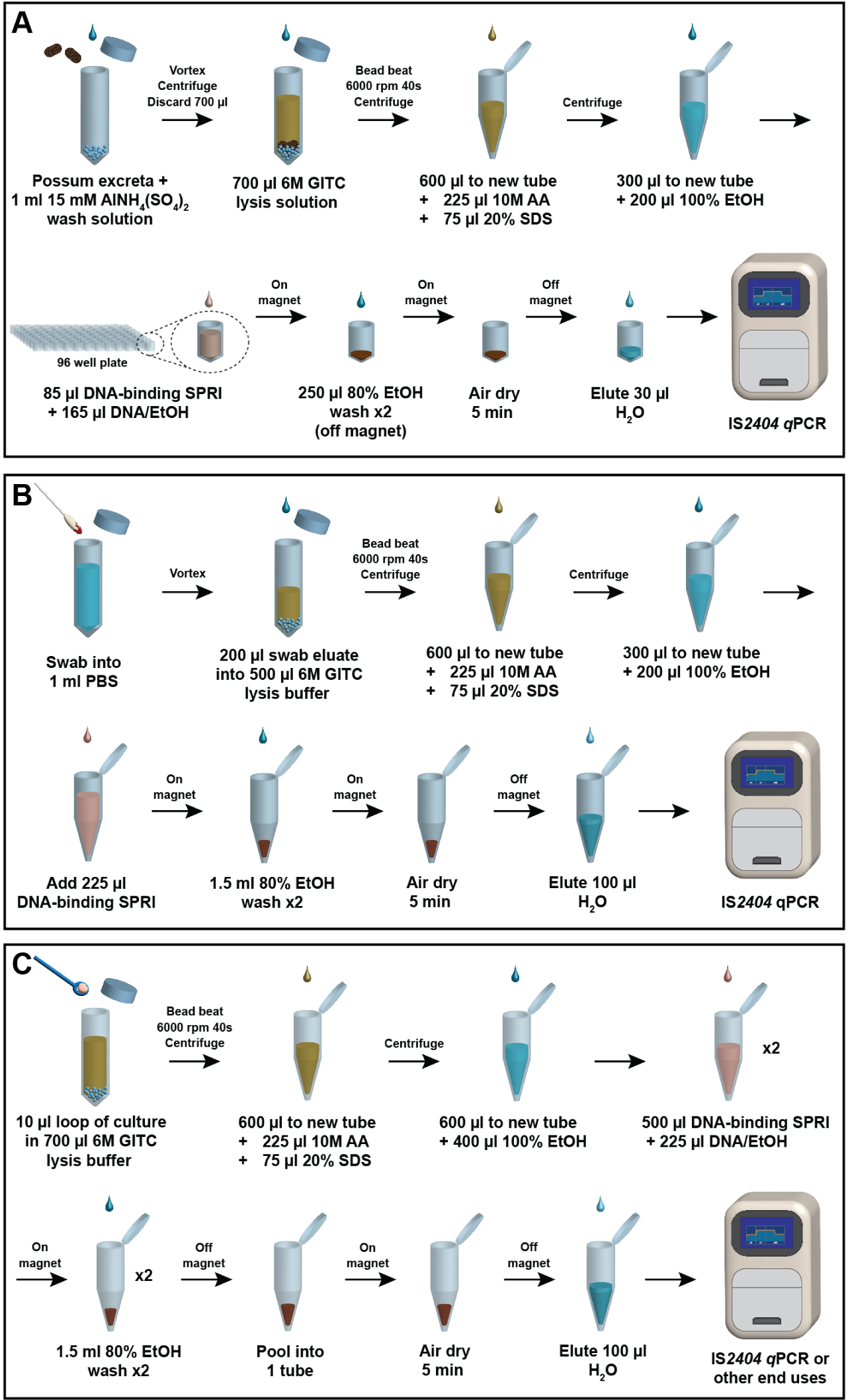
**Overview of solid-phase reversible immobilisation (SPRI)-bead based DNA extraction and purification methods** developed for **A.** Possum excreta specimens; **B.** Biological swabs taken from possums with suspected *M. ulcerans* infection; **C.** Pure mycobacterial culture. GITC: guanidinium thiocyanate, AA: ammonium acetate, SDS: sodium docecyl sulfate, EtOH: ethanol, IS*2404*: insertion sequence *2404*, qPCR: quantitative polymerase chain reaction.

### Mycobacterium ulcerans is stable in possum faecal material

To facilitate development and optimisation of the SPRI-bead DNA extraction method, pooled possum excreta known to be IS*2404* qPCR negative was spiked with a standardised inoculum of *M. ulcerans.* Interval testing of 50 mg of this spiked possum excreta, in triplicate, over 418 days indicated that the concentration of *M. ulcerans* remained stable in this matrix (stored in sealed 50 ml tube at 4oC) over this period, as reflected in unchanged IS*2404* qPCR Ct values (Figure 2A). This pool of stable, spiked excreta was used to create a series of positive possum excreta specimens from 10, 20, 40, 60, 80, 90 to 100 mg for subsequent SPRI-bead assay performance evaluations.

**FIG 2.**
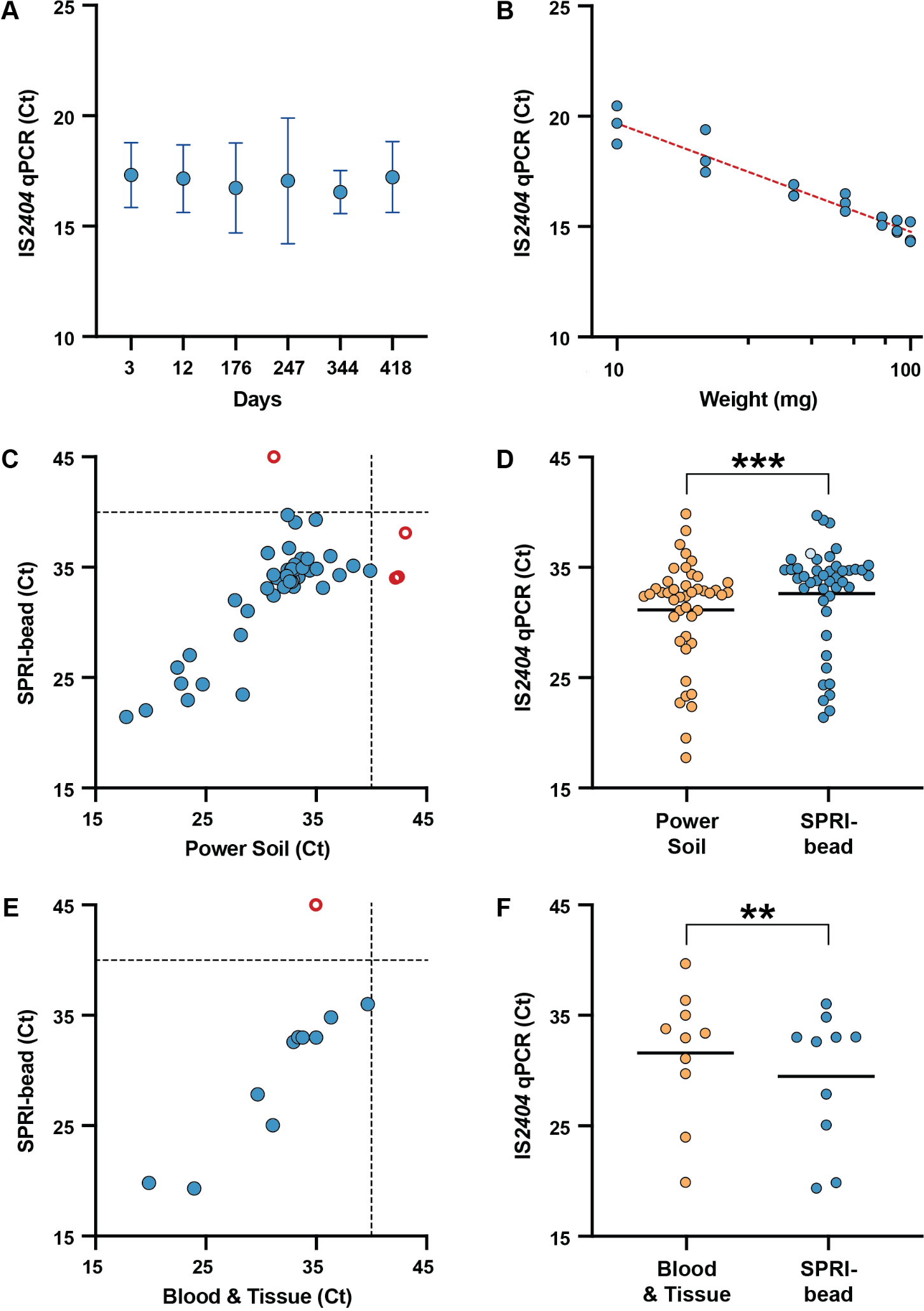
Development and validation of solid-phase reversible immobilisation (SPRI)-bead based DNA extraction methods for detecting *M. ulcerans*. **A.** Stability profile of a contrived *M. ulcerans* positive possum excreta. Blue circles represent the mean of triplicate DNA extractions of 50 mg of the contrived positive sample tested by a quantitative polymerase chain reaction (qPCR) targeting *M. ulcerans* insertion sequence (IS) *2404* at each time point. Error bars represent the 95% confidence interval. **B.** Efficiency of SPRI-bead based DNA extractions of increasing amounts of possum excreta, as assessed by IS*2404* qPCR. Shown are the results of biological triplicate DNA extractions. Dotted red line shows the square of the correlation coefficient, R2 = 0.9237. **C.** Comparison of Qiagen PowerSoil Pro and a SPRI-bead based method for DNA extraction from possum excreta. Possum excreta surveillance samples (N=68) were equally halved and processed by the ‘gold standard’ PowerSoil Pro kit and the SPRI-bead method. Blue circles show 44 samples with DNA extractions that tested IS*2404* positive by both methods. Dotted lines indicate the cycle threshold (Ct) >40 cut-off for the IS*2404* qPCR, above which samples are interpreted as negative. Open red circles show samples not detected by one method. The 20 samples that tested negative by both DNA extraction methods are not shown. **D.** Comparison of Ct values for the 44 possum excreta samples with DNA extractions that tested IS*2404* qPCR positive by both methods. The single pale blue circle represents a SPRI-bead extracted sample that was PCR inhibited on initial testing but tested positive after dilution of the DNA extract. Black horizontal bars represent the mean IS*2404* qPCR Ct value for each method. The null hypothesis (no difference between means) was rejected for p<0.01 (Wilcoxon matched-pairs signed rank test, two-tailed p-value). ***p = 0.0002. **E.** Comparison of the Qiagen Blood & Tissue kit and a SPRI-bead based method for DNA extraction from possum tissue swabs. Swabs collected from possums (N=16) were each eluted in PBS and equal aliquots of eluate were DNA extracted using either the Blood & Tissue Kit or the SPRI-bead method. Blue circles show samples with DNA extractions that tested IS*2404* positive by both methods. The 5 samples that tested negative by both DNA extraction methods are not shown. **F.** The 10 possum swab samples that tested positive by both DNA extraction methods, with Ct values plotted by extraction method. Black horizontal bars represent the mean IS*2404* qPCR Ct value for each DNA extraction method. The null hypothesis (no difference between means) was rejected for p<0.01 (Wilcoxon matched-pairs signed rank test, two-tailed p-value). **p = 0.0020

For quality control, the stability of the 6M GITC lysis solution and the functionalised DNA-binding SPRI beads when used in our assay were also determined. Batch testing of 6M GITC lysis solution of various ages demonstrated consistent performance for at least 1094 days (2 years, 11 months, 29 days; Supplementary Figure 1A). No difference was noted if the solution was stored protected from light at room temperature, then warmed until the precipitated GITC dissolved into solution, or if the 6M GITC buffer was stored protected from light at 28oC throughout. Similarly, batch testing of the DNA-binding SPRI beads performed consistently for at least 1008 days (2 years, 9 months, 2 days; Supplementary Figure 1B). Indicating no deterioration of these solutions over at least 2.8 years.

### Performance of SPRI-bead DNA extraction for possum excreta

DNA was extracted by the SPRI-bead method from triplicate preparations of the seven increasing weights of the *M. ulcerans*-spiked excreta. The resulting DNA preparations were then tested for the presence of *M. ulcerans* by IS*2404*TaqMan qPCR [20]. We observed a consistent, linear reduction in Ct as increasing volumes of possum excreta were tested, up to 100 mg (Fig. 2B). Above 100 mg, PCR inhibition was observed, as monitored by presence or absence of the qPCR internal positive control (Table S1). This observed decrease in performance when processing above 100 mg of excreta was likely due to the higher quantity of humic acids and other interferents present in possum excreta that were not effectively removed during extraction.

### Comparison of SPRI-bead DNA extraction with DNeasy PowerSoil Pro kit for possum excreta

Having established the maximum volume of material the SPRI-bead DNA extraction method could process, we sought to further validate the protocol by screening 68 possum excreta specimens from a previous study we knew to have originated from sites with a high probability of harbouring *M. ulcerans* [8]. Individual excreta pellets were divided into equal halves (determined by weight), with half processed using our SPRI-bead method and the other half the DNeasy PowerSoil Pro Kit (considered the ‘gold-standard’). For large brushtail possum excreta pellets which could exceed 300 mg in weight, 50 mg of the excreta were tested by each extraction method. The weights of possum excreta tested ranged from 21 mg to 96 mg, with an average weight of 53 mg (95% confidence interval 50 – 55 mg) (Supplementary Figure 2). The resulting DNA preparations were tested for the presence of *M. ulcerans* using the IS*2404* qPCR. A comparison of the efficacy of the two DNA extraction methods when processing the paired possum excreta samples is shown in Figure 2C and D. Of the 68 possum excreta samples extracted: 44 were IS*2404* PCR positive by both methods, 20 were IS*2404* PCR negative by both methods, while 4 samples were detected by only one method (Figure 2C).

Three samples had discordant IS*2404* qPCR results, testing positive when extracted with the SPRI-bead method with Ct values ranging from 34 – 38, but negative when extracted by the gold standard method with corresponding sample Ct values of 42 – 43. These three samples were interpreted as negative due to being above the positive interpretation cut-off for the IS*2404* qPCR of Ct <=40. PowerSoil Pro extractions of additional possum excreta collected from the same locations at the same time as each of the three discordant samples had IS*2404* Ct values of 32 – 34, suggesting these were likely true positive samples. Two samples that were IS*2404* positive when extracted by the gold standard method initially yielded “no result” following SPRI-bead extraction due to PCR inhibition. Testing 1/2 and 1/4 dilutions of the SPRI-bead DNA extracts enabled one of these samples to be detected at both dilutions (1/2 dilution: Ct 37.69, 1/4 dilution: Ct 36.30), however the other sample remained inhibited at both dilutions and was interpreted as a false negative result. This PCR inhibited sample was 96 mg in weight. Based on the above, there were 48 true positive and 20 true negative samples. Therefore, our in-house DNA extraction method had a sensitivity of 97.9%, specificity of 100.0%, positive predictive value of 100.0% and negative predictive value of 95.2%.

For the 44 samples from which *M. ulcerans* was detected in the DNA extractions processed by both methods, there was overall good concordance between the two methods (Spearman r=0.6947, p<0.0001). However, the mean IS*2404* Ct of 32.65 obtained with the SPRI-bead method was significantly higher than the PowerSoil Pro mean Ct of 31.18 (p=0.0002) (Fig. 2D). This mean difference of 1.47 qPCR cycles, suggested an approximate 3-fold decrease in sensitivity of the SPRI-bead method compared to the PowerSoil Pro kit.

### Comparison of SPRI-bead DNA extraction with DNeasy Blood & Tissue kit for swabs taken from possums

We also explored the suitability of the SPRI-bead method for detecting *M. ulcerans* DNA from swabs collected from live or necropsied possums, some with ulcerated lesions suspicious of *M. ulcerans* infection. A total of 16 swabs, collected from 10 possums were available for testing (see Supplementary Table 2 for details of possum specimens) and stored at -20oC until processed. Swab material was eluted in PBS before equal volumes of the eluate were processed for DNA extraction using the DNeasy Blood & Tissue Kit (considered the ‘gold-standard’ for comparison purposes) and a SPRI-bead method omitting the alum wash step (outlined in Figure 1B). The DNA extraction from each method was then tested for the presence of *M. ulcerans* by IS*2404* qPCR. Using the DNeasy Blood & Tissue kit, 11 swab eluates tested positive and 5 were negative, compared with 10 positive and 6 negative swab eluates using the SPRI-bead method. The single false-negative result from the SPRI-bead method corresponded with an eluate that had been stored for >300 days before testing by this method and had a relatively high Ct value of 35, therefore this specimen may have degraded during storage beyond the limit-of-detection for the IS*2404* qPCR. Based on this small sample size, the sensitivity of the SPRI-based method was 90.9%, specificity 100.0%, positive predictive value 100.0%, and negative predictive value 83.3%.

Similar to the possum excreta DNA extracts, there was good concordance between the two methods (Spearman r=0.9362, p=0.0002; Figure 2E). However, in this comparison, the SPRI-bead method was observed to have a significantly lower mean Ct of 29.44 than the mean Ct of 31.56 for Blood & Tissue Kit (p=0.0020; Figure 2F). The mean difference of 2.12 PCR cycles indicated the SPRI-bead method was approximately 4-fold more sensitive than the gold standard method.

### SPRI-bead DNA extraction for mycobacterial cultures

We also adapted the SPRI-bead method for preparation of genomic DNA from pure cultures of *M. ulcerans* and other mycobacteria (Figure 1C). The method omits the alum wash step since humic substances are not present in pure bacterial culture, and an increased volume of lysate is processed (1000 µl compared to 165 µl) to maximise DNA yield. Three genomic extractions of *M. ulcerans* yielded an average of 24.2 µg (range 20.8 – 29.6 µg) of DNA with a peak size of 8,865 bp (range 6,822 to 10,314 bp). Similarly high amounts of genomic DNA were also obtained from *Mycobacterium abscessus* (23.3 µg; 10,336 bp), *Mycobacterium chimera* (14.5 µg; 10,910 bp), *Mycobacterium marinum* (26.8 µg; 12,586 bp), *Mycobacterium smegmatis* (28.3 µg; 10,657 bp) and *Mycobacterium virginiense* (28.4 µg; 13,619 bp). Full quality control metrics of these mycobacterial genomic DNA extractions and two comparators, including a chloroform extraction of *M. ulcerans* (10,462 bp) [41], and a high quality *Staphylococcus epidermidis* extraction performed using the Qiagen Blood & Tissue kit with pre-treatment to weaken the gram positive cell wall [42] that was subsequently PacBio sequenced [43] (56,896 bp), is shown in Supplementary Figure 3.

### Cost comparison of DNA extraction methods

We assessed the material costs needed to prepare DNA from 96 excreta samples by both our SPRI-bead method and Qiagen PowerSoil Pro kit. The cost per sample for the PowerSoil Pro method depended on kit size, with a kit of four 96-well plates costing AUD $8 per sample, increasing to AUD $12 per sample for the smaller, 50-tube kit. By comparison, our SPRI-bead method cost AUD $2 per sample irrespective of 396 or 50 samples being processed. Similarly, the cost for processing biological swabs collected from possums using the Qiagen Blood & Tissue kit was AUD $10 per sample for a 50-tube kit, whereas our SPRI-bead method was only AUD $2.

## DISCUSSION

Extensive environmental surveillance studies have been the foundation of field-based research leading to the discovery that BU is a zoonosis [8, 9, 10, 11, 12]. Possum excreta surveys have been instrumental in ascertaining transmission pathways and predicting the geographic risk of BU spread to humans in Victoria, Australia [8, 10]. Molecular analytical methods based on IS*2404* qPCR have been the mainstay for these surveys, but the high sensitivity and specificity of this assay [20] is countered by the relatively high cost of the commercial nucleic acid extraction reagents needed. For environmental surveillance projects involving the collection of thousands of excreta specimens, research budgets can be quickly exhausted, hindering endeavours to halt disease spread through large-scale measure-intervene-measure public health responses. Here, we addressed this bottleneck by developing lower-cost nucleic extraction methods.

Integral to all three of our described methods were the 6M GITC lysis solution and functionalised DNA-binding SPRI beads. When stored appropriately, we have demonstrated both reagents perform consistently for at least 2.8 years, indicating they are amenable to bulk preparation for later use, removing the onus of preparing multiple solutions immediately prior to testing. Furthermore, use of in-house solutions overcomes supply chain issues present with commercial reagents and kits, such as the global dearth of nucleic acid extractions kits experienced during the SARS-CoV-2 pandemic [44, 45].

In view of the large number of possum excreta routinely processed in *M. ulcerans* surveillance studies, we prioritised a protocol amenable to high throughput processing of samples in a 96-well plate format rather than extraction of the maximal volume of lysate in individual tubes. Therefore, we down scaled our method to process 1/9 of the lysed possum excreta sample (300 µl of 900 µl lysate post protein precipitation, then 165 µl of the 500 µl of precipitated DNA), as compared to the PowerSoil Pro kit which used the full lysate volume. Despite the much lower fraction of sample extracted, paired comparison of extractions indicated our SPRI-bead method was only approximately three-fold less sensitive than the commercial kit, with overall good concordance in results (r=0.6947, p<0.0001), and a calculated sensitivity of 97.9% and specificity of 100.0% for the 68 possum excreta samples tested.

Discordance between the two methods for three of the possum excreta samples may have been due to unequal distribution of *M. ulcerans* within the excreta specimens, resulting in failure of the PowerSoil Pro method to detect the known positive samples. Of note, the single SPRI-bead extracted sample that failed to yield a result due to PCR inhibition, corresponded with the heaviest surveillance specimen tested (96 mg), that was close to the 100 mg maximum extractable weight determined when developing the method based on a contrived *M. ulcerans* spiked possum excreta. Due to dietary differences between individual possums, the humic substance content of excreta samples can vary. This PCR-inhibited result suggested that for some possum excreta samples even 100 mg may contain humic compounds that exceed the flocculation capacity of the 15 mM AlNH_4_(SO4)_2_ wash, therefore extraction of a lower weight of these samples is recommended if dilution of the DNA extraction is insufficient to overcome PCR inhibition. It could also be argued that routine use of a lower sample input maybe prudent to avoid PCR inhibition, without necessarily incurring a high number of false negatives results. Indeed, the average weight of the possum excreta samples tested by each method in the experiment was only 53 mg. Therefore, we recommend using between 50 to 90 mg of possum excreta when performing DNA extraction with our SPRI-bead method.

Since fewer possum swabs are collected during surveillance studies compared to possum excreta samples, for this DNA extraction method we developed a tube-based method in which 1/3 of the lysed sample was extracted. Although pairwise comparison of the possum swab samples demonstrated our SPRI-bead method was superior, with an estimated four-fold sensitivity above the commercial Blood & Tissue kit, calculation of the overall sensitivity of the method was hindered by the small experimental sample size of 16 swabs and availability of only retrospective samples for testing (around which our method was developed). Since only 11 true positive samples were tested, the single false negative resulted in a calculated sensitivity of only 90.9%, with a specificity of 100.0%. Of note, this false negative sample was from sample with a relatively high a Ct of 35 when tested by the gold standard Blood & Tissue Kit, that had been stored for over 300 days prior to testing by the SPRI-bead method, therefore this specimen may have degraded during storage beyond the limit-of-detection for the IS*2404* qPCR. The sensitivity testing of our method was disadvantaged by being limited to retrospective testing of existing samples that were eluted in 1 ml of PBS then stored at -20oC for months after an aliquot was tested by the gold standard method. Ideally, if prospective testing were performed, swabs would be resuspended directly in the 6M GITC solution immediately after collection, minimising the dilution of the sample in PBS and deterioration of DNA over time and from repeated freeze thawing.

Our SPRI-bead based methods yielded DNA extractions from environmental samples, diagnostic swabs and pure mycobacterial cultures that were suitable for end use in IS*2404* PCR reactions. To functionalise the DNA-binding SPRI beads we used the method described by Jolivet & Foley, which was developed as a low-cost substitute for AMPureXP (Beckman Coulter) beads, designed for DNA clean-up [37]. While the IS*2404* element is around 1,300 bp in size [20, 25], the product of the IS*2404* qPCR used is only 59 bp [20], and the original study validating the IS*2404* PCR method used amplicons that were 515 bp in length as a positive control [20]. Our protocol used a SPRI:DNA ratio of 0.5 to target approximately 600 bp, rather than the 1.8 times ratio typically recommended by most commercially available SPRI-bead based DNA clean-up reagents that target a peak of 200 bp, optimised for PCR and next generation sequencing [46]. This partially accounts for the relatively low average peak DNA size of our mycobacterial genomic DNA extractions (10,588 bp), as compared to the commercial kit extraction of *S. epidermidis* (56,896 bp). While bead beating can result in DNA shearing, the upper size of >60,000 bp for the *M. marinum*, and 52,397 bp for the *M. virginiense* genomic extractions using our SPRI-bead method, indicates the method is capable of extracting long length DNA fragments. However, if DNA is specifically intended for long read sequencing, the use of SPRI-beads functionalised for high molecular weight DNA selection (through alteration of polyethylene glycol (PEG) 8000 and NaCl concentrations [47]) or a commercial product like SPRIselect (Beckman Coulter) would be recommended instead, with inclusion of an RNA depletion step.

Interspecies differences also contribute to the lower average peak DNA size for our mycobacterial genomic extractions compared to the *S. epidermidis* isolate. The mycobacterial cell wall is rich in lipids and polysaccharides forming a complex structure that renders the bacterium resistant to lysis, hindering most conventional DNA extraction methods [48, 49], therefore most protocols developed for mycobacteria are laborious (potentially taking days), with many requiring phenol or phenol/chloroform [49]. A chloroform-based DNA extraction method [41] included as a comparator, used for the genomic DNA extraction of *M. ulcerans* JKD6049 had an average peak DNA size of only 10,462 bp, highlighting the difficulties in obtaining high quality, long-length mycobacterial DNA.

Here we present low-cost, SPRI-bead based methods for the extraction of *M. ulcerans* from Australian possum excreta and swabs from possums with suspected *M. ulcerans* infection, as well as the extraction of mycobacterial genomic DNA from pure cultures, which demonstrate comparable performance to commercially available kits, at around 1/6 to 1/5 the cost. Although developed for the IS*2404* PCR detection of *M. ulcerans* from possum excreta, swabs from possums and pure bacterial culture, the described methods should be adaptable for the testing of *M. ulcerans* from other environmental sources and swabs from animals other than possums, as well as other bacteria of interest when paired with an appropriate diagnostic PCR. While not implemented in this study, we have previously demonstrated SPRI-bead based nucleic acid extractions to be amenable to high-throughput, low-cost, automation in a 96-well format [36], which could be an area for future assay development.

Collectively, over 4,700 possum excreta samples were tested in the four Victorian *M. ulcerans* surveillance studies discussed in the introduction [8, 9, 11, 12]. The 2023 study by Vandelannoote et al., contributed 2,282 samples [8], with over AUD $20,000 spent on commercial DNA extraction kits for this single study. When performed on such scale, the expenditure required for sample processing can be prohibitive to ongoing surveillance when reliant on high-cost commercial reagents. A means to circumvent this barrier prompted this study, and as demonstrated, has been achieved.

The cumulative knowledge from two decades of *M. ulcerans* environmental surveys in Victoria now enables us to predict areas in which humans are at increased risk of disease acquisition and provides local government with a tangible means of interrupting transmission pathways through targeted mosquito control programs [50]. However, successful implementation of such a program requires ongoing possum excreta surveillance to inform prediction models and is subject to resourcing and budgetary constraints. By providing an economical means to continue *M. ulcerans* environmental surveillance surveys, this study aims to facilitate efforts to halt and reverse the spread of human BU disease in Victoria. Should these endeavours prove successful, they will also provide vital insights into means to disrupt the spread of BU in other endemic regions in the world.

## ETHICS APPROVAL

Opportunistic native possum necropsy-based and active trap-and-release surveillance studies were conducted under ethical approvals from the University of Melbourne AEC 22910 and DELWP permits 10009447 and 10010257.

