## Supplementary data for "A low-cost and versatile paramagnetic bead DNA extraction method for *Mycobacterium ulcerans* environmental surveillance"

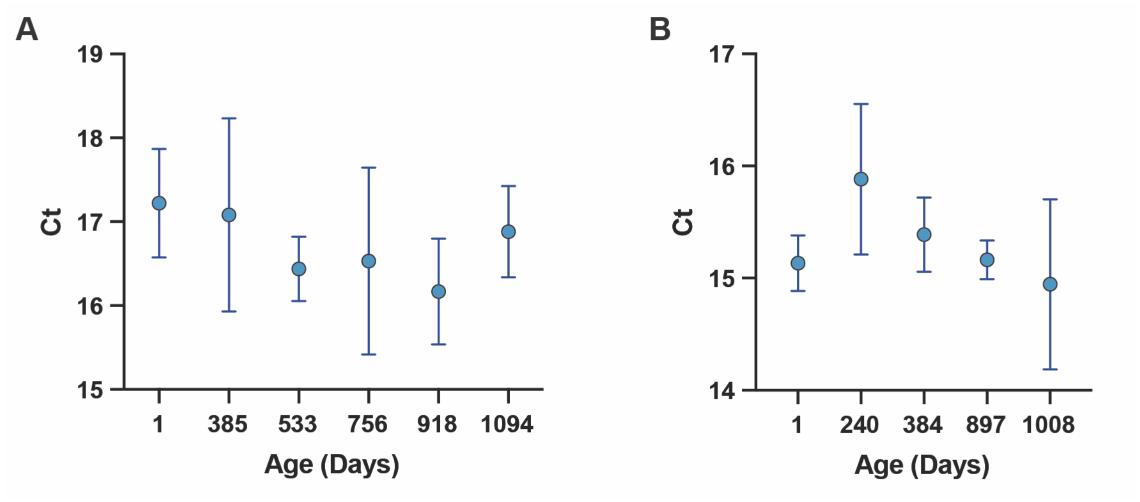


**Supplementary Figure 1. Quality control for 6M guanidinium thiocyanate (GITC) lysis solution and functionalised DNA-binding solid-phase reversible immobilisation (SPRI) beads. A.** Batches of 6M GITC lysis solution of varying ages were used in the extraction of 50 mg of contrived positive, *M. ulcerans* containing possum excreta, the resultant DNA extraction was then assayed with IS*2404* PCR. Each batch was tested in triplicate. Testing of all batches was performed concurrently. Plotted points represent the mean for each replicate, error bars indicate the 95% confidence interval. There was no significant difference between the lysis ability of each batch [Null hypothesis (no difference between means) was accepted for $P < 0.05$. Assessed using one-way analysis of variance (ANOVA) with Tukey's test for multiple comparisons]. **B.** Batches of DNA-binding SPRI beads of varying ages were used in the extraction of 50 mg of contrived positive, *M. ulcerans* containing possum excreta, the resultant DNA extraction was then assayed with IS*2404* PCR. Each batch was tested in triplicate. Testing of all batches was performed concurrently. Plotted points represent the mean for each replicate, error bars indicate the 95% confidence interval. There was no significant difference in between the DNA-binding of each batch [Null hypothesis (no difference between means) was accepted for $P < 0.05$. Assessed using one-way analysis of variance (ANOVA) with Turkey's test for multiple comparisons].

**Supplementary Table 1. Weight series of contrived *M. ulcerans*-positive possum excreta.** A weight series of contrived *M. ulcerans*-positive possum excreta was DNA extracted using a solid-phase reversible immobilisation (SPRI)-bead based method then tested for the presence of *M. ulcerans* using a TaqMan quantitative polymerase chain reaction (qPCR) targeting insertion sequence (IS) *2404* detected using a FAM labelled probe. The IS*2404* qPCR is Multiplexed with an exogenous internal positive control (IPC) that uses a VIC probe, which assists in distinguishing a true negative reaction from one that has no result due to PCR inhibition.

| **Weight (mg)** | **Replicate 1** | | **Replicate 2** | | **Replicate 3** | |
|---|---|---|---|---|---|---|
| | IS*2404* qPCR (Ct) | IPC (Ct) | IS*2404* qPCR (Ct) | IPC (Ct) | IS*2404* qPCR (Ct) | IPC (Ct) |
| 10 | 20.464 | 26.337 | 19.681 | 26.484 | 18.747 | 26.262 |
| 20 | 19.393 | 26.339 | 17.473 | 26.194 | 17.974 | 26.269 |
| 40 | 16.912 | 25.994 | 16.409 | 26.371 | 16.389 | 26.354 |
| 60 | 16.497 | 26.234 | 16.066 | 26.495 | 15.697 | 26.332 |
| 80 | 15.406 | 27.224 | 15.437 | 27.93 | 15.06 | 26.551 |
| 90 | 15.283 | 26.084 | 14.741 | 24.805 | 14.808 | 26.396 |
| 100 | 14.383 | 26.016 | 14.315 | 26.772 | 15.218 | 26.142 |
| 110 | Undetermined | Undetermined | Undetermined | Undetermined | Undetermined | Undetermined |

Ct: cycle threshold.

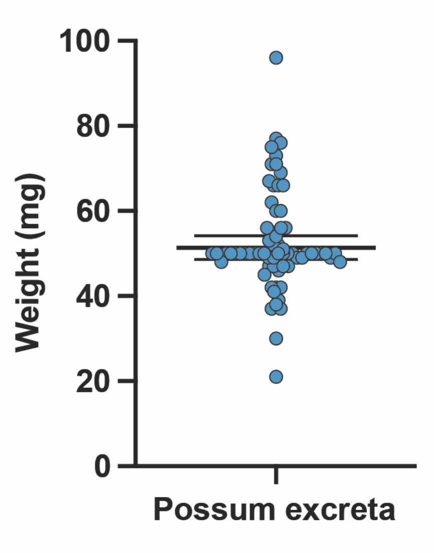


**Supplementary Figure 2. Weights of possum excreta tested for validation of a SPRI-bead based DNA extraction method.** Sixty-eight possum excreta surveillance samples were divided into two equal halves (by weight) with one half processed by the 'gold standard' PowerSoil Pro kit, the other by our SPRI-bead method. For large excreta pellets, where half the pellet would exceed the predicated 100 mg maximum input for the SPRI-bead method, 50 mg of the pellet was tested by each extraction method. Each circle represents the weight of excreta processed by both DNA extraction method, for each of 68 samples. Thick horizontal line represents the median weight, error bars indicate the 95% confidence interval.

**Supplementary Table 2. Comparison of the efficiency of a SPRI-bead method with the DNeasy Blood & Tissue kit for extracting *M. ulcerans* DNA from swabs sampled from Australian possums as determined by IS*2404* qPCR.**

| Possum number (ID) | Swabbed site | Blood & Tissue (Ct) | SPRI-bead (Ct) |
|---|---|---|---|
| 1 (W379-23) | Multiple ulcers | ND | ND |
| 2 (W381-23) | Multiple ulcers | ND | ND |
| 3 (W386-23) | Multiple ulcers | ND | ND |
| 4 (W387-23) | Multiple ulcers | ND | ND |
| 5 (W380-23) | Multiple ulcers | 19.836 | 19.814 |
| 6 (6/22) | Oral cavity | 36.346 | 34.806 |
| | Cloaca | 29.675 | 27.849 |
| 7 (13/22) | Oral cavity | 39.652 | 36.017 |
| | Cloaca | ND | ND |
| 8 (14/22) | Oral cavity | 31.072 | 25.043 |
| | Cloaca | 32.946 | 32.578 |
| | Pouch | 33.367 | 33.022 |
| 9 (15/22) | Oral cavity | 34.952 | ND |
| 10 (20/23) | Oral cavity | 33.762 | 32.982 |
| | Cloaca | 34.979 | 32.982 |
| | Ulcer (L hind paw) | 23.931 | 19.301 |

Ct: cycle threshold, ND: not detected, L: left.

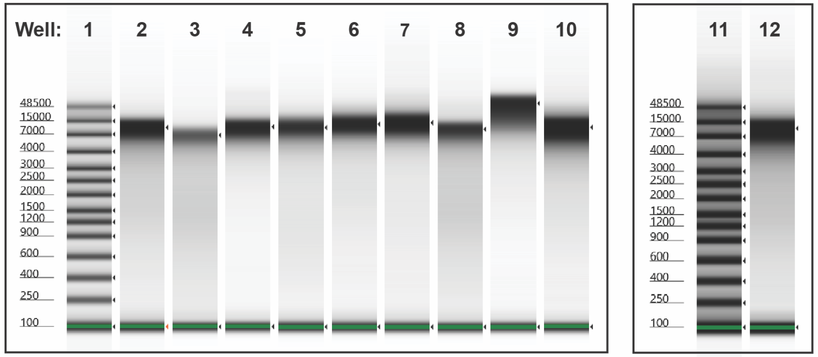


| Well | Organism | Strain | Extraction method | Peak (bp) | % Integrated area | From (bp) | To (bp) | Isolate Reference |
|---|---|---|---|---|---|---|---|---|
| 2 | *M. ulcerans* | JKD8049 | SPRI-bead | 10,314 | 96.57 | 3,808 | 25,279 | (1) |
| 3 | *M. ulcerans* | JKD8049 | SPRI-bead | 6,822 | 92.33 | 3,553 | 16,669 | (1) |
| 8 | *M. ulcerans* | JKD8049 | SPRI-bead | 9,458 | 97.32 | 3,862 | 22,130 | (1) |
| 5 | *M. abscessus* | TPS8830 | SPRI-bead | 10,336 | 96.15 | 3,940 | 30,295 | Unpublished |
| 12 | *M. chimera* | DMG1600132 | SPRI-bead | 10,910 | 91,59 | 4,236 | 28,628 | (2) |
| 6 | *M. marinum* | "M" strain | SPRI-bead | 12,586<br>>60,000 | 87.03<br>0.89 | 4,865<br>>60,000 | 54,651<br>>60,000 | (3) |
| 4 | *M. smegmatis* | MC2155 | SPRI-bead | 10,657 | 96.77 | 4,087 | 25,549 | (4) |
| 7 | *M. virginiense* | TPS8833 | SPRI-bead | 13,619 | 96.25 | 4,850 | 52,397 | Unpublished |
| 10 | *M. ulcerans* | JKD8049 | Chloroform extraction | 10,462 | 99.30 | 3,920 | 37,734 | (1) |
| 9 | *S. epidermidis* | BPH0747 | Qiagen Blood & Tissue kit | 56,896 | 99.01 | 6,063 | >60,000 | (5) |

**Supplementary Figure 3. Comparison of DNA extractions from pure mycobacterial cultures using SPRI-bead based method.** Note: non-standardised quantity of DNA loaded per well.
